# Dietary acculturation of Asian and the Middle East and North African region immigrants to Saudi Arabia: healthy or unhealthy acquired nutritional behavior?

**DOI:** 10.1101/591230

**Authors:** Rasmieh Al Zeidan, Shabana Tharkar, Ahmed Hersi, Anhar Ullah

## Abstract

Travel and migration influences food behavior. This study assessed the dietary acculturation of immigrants in Saudi Arabia, with regard to length of stay and health status of immigrants. This cross-sectional study included 880 university immigrant employees and their family members from Asian and Middle East and North African regions. Dietary acculturation was assessed based on knowledge and practice of methods of food preparation, type of food consumption, and nutrition label reading behavior, using a measurement tool on a 5-point Likert scale. Furthermore, a scoring system was adapted for healthy practices. Anthropometric, biochemical, and blood pressure measurements were performed as per the World Health Organization guidelines, to determine participants’ health and comorbid status. In addition, scores were calculated for healthy options. Factors influencing better awareness were determined by logistic regression analysis. The most adopted methods of food preparation after migration were barbeque (p=0.018), microwave cooking (p=0.002), and raw food consumption (salads) (p<0.001). Consumption of carbonated drinks (p=0.025), fried fatty and processed food (p=0.037), and sweets and candies (p=0.008) were significantly higher among recent immigrants of <5 years of residency. Label reading behavior of nutritional contents and low-fat options was higher among immigrants with ≥5 years duration of residency (63%; p<0.001). Although female gender, longer duration of residency in Saudi Arabia and presence of comorbidity significantly improved the overall awareness and practice scores in the binary analysis, they failed to show significance in regression model except for the presence of diabetes which improved only awareness. None of the other independent factors seem to influence healthy practices. Chronic diseases like obesity, diabetes and hypertension increased with longer duration of migration(p<0.001).

New immigrants are at risk of acquiring negative dietary habits compromising health, necessitating follow-up studies to establish causation. Interventional policy measures are recommended to formulate dietary guidelines.

## Introduction

Immense technological developments and globalization has led to the rapid rise in migration in the 21^st^ century, enhancing cross-culture perceptions, especially in food and lifestyle. International statistics on migration show that Asian countries are in the lead among those targeting the West as a preferred destination [1]. The world has also witnessed a concomitant rise in prevalence of lifestyle-related disorders like overweight and obesity, diabetes, hypertension, and other cardiovascular diseases [2]. There is evidence establishing significant link between adaptation to the new culture and non-communicable diseases [3]. This suggests the need to explore the adaptive changes of immigrants by the process of acculturation, “the process by which the migrant group adopts the cultural pattern of the dominant group” [4]. The culturally diverse environment offers a favorable atmosphere for the adoption of host country lifestyle, by gradually relinquishing the native habits. Whereas some studies found positive effects of dietary acculturation (like increased consumption of fruits and vegetables and lowered intake of fried-food), others documented negative effects (like increased intake of carbonated drinks and fatty and processed food), thus compromising health [5-9]. Although, most countries demand strict regulations on health as a prerequisite condition for immigration, several studies have reported the rapid deteriorating in health status of immigrants by the rising prevalence of comorbidities (like dyslipidemia, metabolic syndrome, and other cardiovascular disorders), over time [10-12]. Saudi Arabia is an oil rich kingdom with a gross domestic product of 21,120 USD in 2017 with a likely projected increase of 2% in 2019, as stated by the World Bank [13]. The Kingdom offers a huge international job market with nearly 33% of the employees being immigrants [14]. The low cost and high affordability of abundant multi-cuisine food poses a risk towards increased consumption of high-calorie food, processed food and sweetened drinks, thus jeopardizing health. A cohort study found an association between risk factors such as fast-food consumption and decreased intake of fruits and vegetables with cardiac events among immigrant employees of a large university [15]. Despite the large size of the immigrant population in the Kingdom of Saudi Arabia, literature documenting the influence of acculturation on the immigrants’ dietary practices and food behavior does not exist. Therefore, the present study aims to investigate changes in food behavior of immigrants, in relation to the duration of residency and presence of comorbidities. An insight into the nutritional behavior of immigrants may indirectly reflect on the country’s dietary pattern. The data might supplement necessary information for future health promotion and prevention programs for the population by policy makers.

## Methods

### Study design, setting, and participants

Heart Health Promotion (HHP) is a prospective registry enrolling 4500 university employees and their family members from the largest and top-ranking institution in Saudi Arabia. The university offers courses in the fields of medicine, engineering, natural sciences, and humanities. Of the total 1437 immigrants from Middle East and North African (MENA) and Asian region that were invited, 60% (n=880) agreed to participate in the sub-study, a cross-sectional study. The objective of the study was to assess the dietary habits of immigrants before and after migration. Participants were from Asian countries (India, Pakistan, Sri Lanka, and Bangladesh) and the MENA region (Syria, Iraq, Lebanon, Egypt, Sudan, Kuwait, Jordan, and Palestine).

### Ethical considerations

The initial study was approved by the institutional review Board (IRB) of the University (reference number 13–3721). The study was conducted in accordance with the guidelines of Helsinki Declaration. The study participants were informed of the purpose of the study and all the procedures involved in completing the questionnaire and obtaining the clinical measurements, before written consents were obtained. Anonymity was maintained to preserve confidentiality.

### Data collection

The present study used two questionnaires for data collection. The first tool measured dietary changes and acculturation on a 5-point Likert scale while the second tool was adapted from the WHO-STEPwise approach to chronic disease risk factor surveillance (STEPS), for anthropometric and biochemical measurements. Data collection was done in the first quarter of 2014 by well-trained interviewers who administered the questionnaire to the participants.

#### Description of questionnaire

Dietary acculturation was assessed using a validated tool adapted from Chinese immigrants’ study by Rosenmöller et al [8]. The questionnaire addressed perceived changes in dietary practices in terms of food preparation, dietary pattern, and nutritional knowledge and awareness, since the participants’ migration to Saudi Arabia. Questions to assess changes in dietary pattern, and knowledge and awareness regarding nutrition and dietary behavior are shown below:

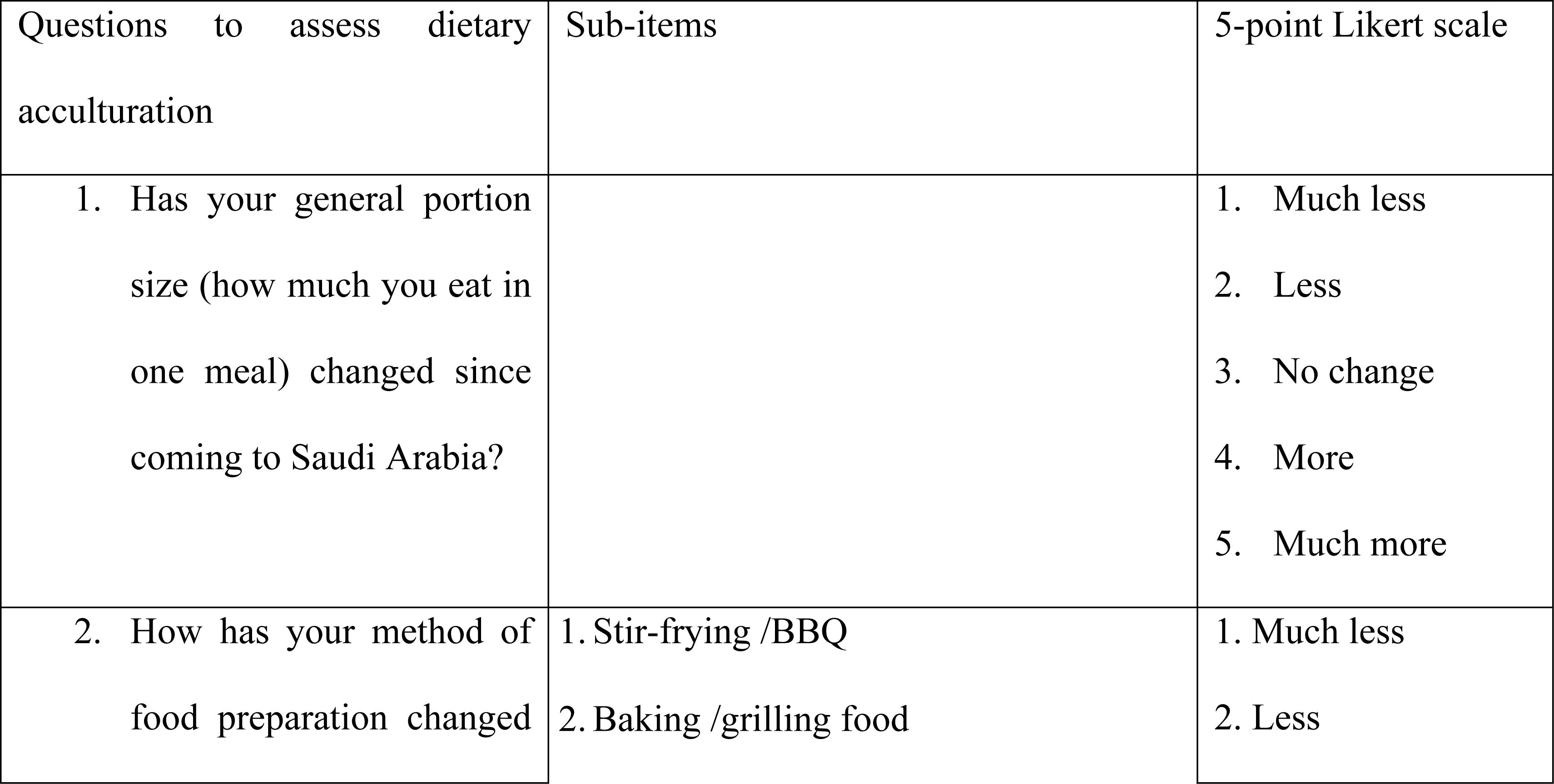
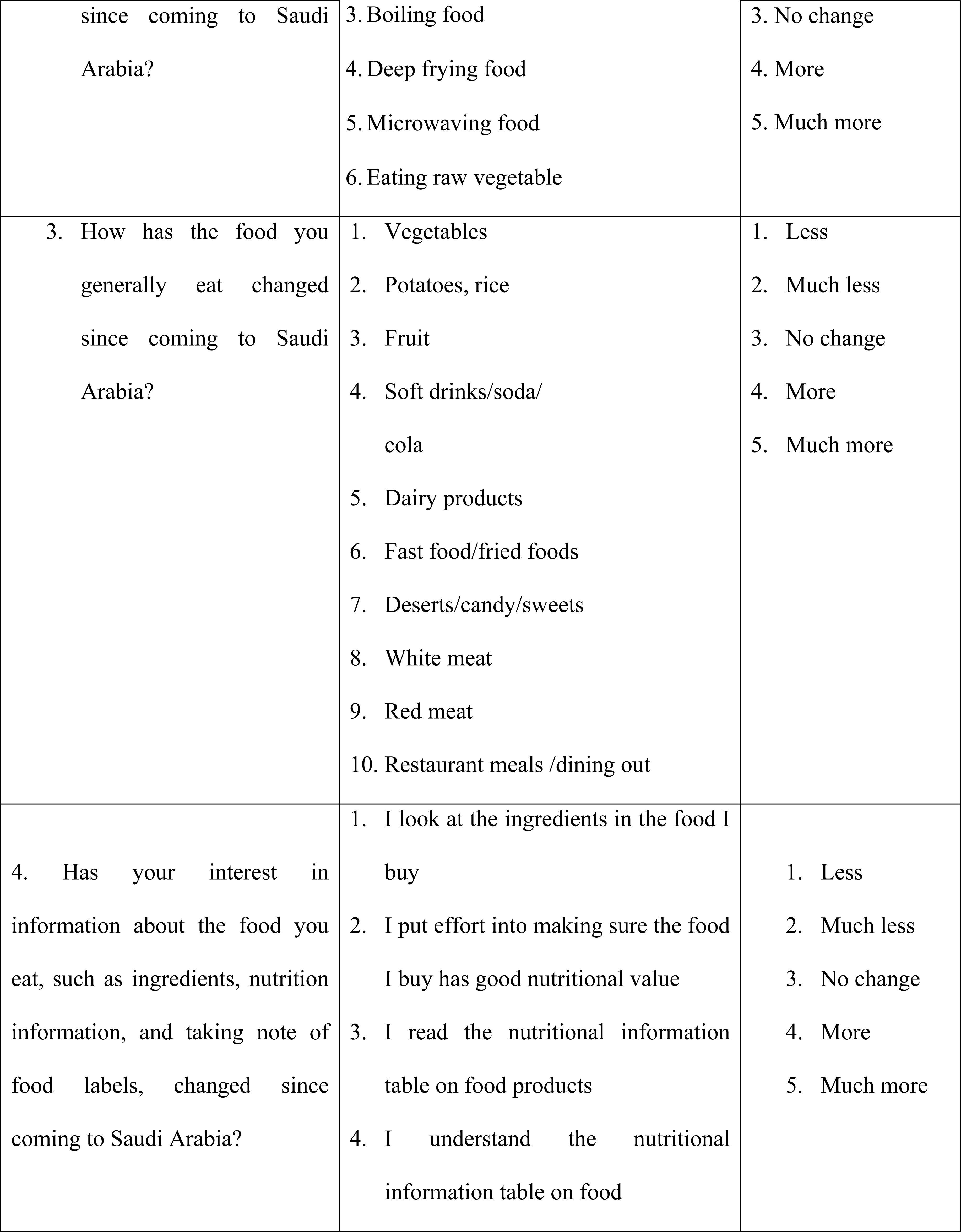
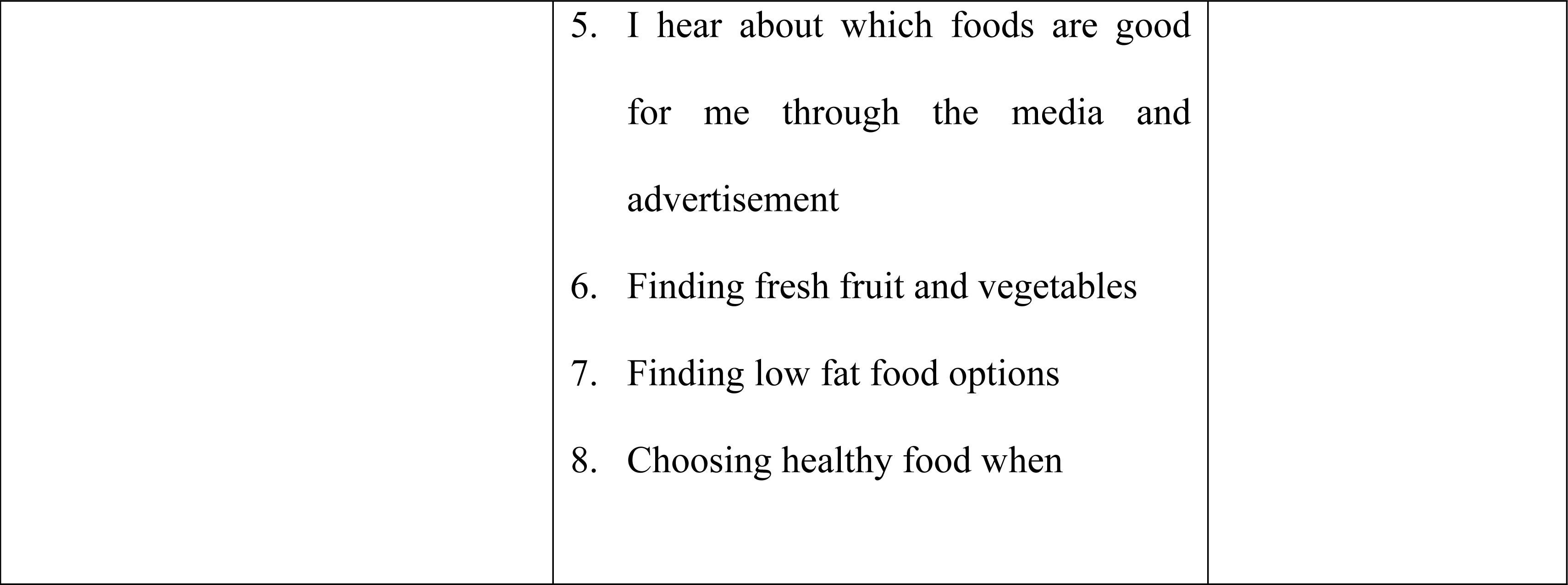

| Questions to assess dietary acculturation | Sub-items | 5-point Likert scale |
| --- | --- | --- |
| 1. Has your general portion size (how much you eat in one meal) changed since coming to Saudi Arabia? |  | 1. Much less<br>2. Less<br>3. No change<br>4. More<br>5. Much more |
| 2. How has your method of food preparation changed | 1. Stir-frying /BBQ<br>2. Baking /grilling food | 1. Much less<br>2. Less |
| since coming to Saudi Arabia? | 3. Boiling food<br>4. Deep frying food<br>5. Microwaving food<br>6. Eating raw vegetable | 3. No change<br>4. More<br>5. Much more |
| 3. How has the food you generally eat changed since coming to Saudi Arabia? | 1. Vegetables<br>2. Potatoes, rice<br>3. Fruit<br>4. Soft drinks/soda/cola<br>5. Dairy products<br>6. Fast food/fried foods<br>7. Deserts/candy/sweets<br>8. White meat<br>9. Red meat<br>10. Restaurant meals /dining out | 1. Less<br>2. Much less<br>3. No change<br>4. More<br>5. Much more |
| 4. Has your interest in information about the food you eat, such as ingredients, nutrition information, and taking note of food labels, changed since coming to Saudi Arabia? | 1. I look at the ingredients in the food I buy<br>2. I put effort into making sure the food I buy has good nutritional value<br>3. I read the nutritional information table on food products<br>4. I understand the nutritional information table on food | 1. Less<br>2. Much less<br>3. No change<br>4. More<br>5. Much more |
|  | 5. I hear about which foods are good<br>for me through the media and<br>advertisement<br>6. Finding fresh fruit and vegetables<br>7. Finding low fat food options<br>8. Choosing healthy food when |  |

### Measurements

The second questionnaire is a modified form of WHO-STEPS (Arabic and English Forms) with the following sequential data collection steps:

Step I – demographic questions; II-anthropometric measurements; and III-biochemical measurements [16].

#### Anthropometric measurements

These included height, weight, and waist circumference. Body mass index (BMI) was then calculated as the ratio of weight in kilograms and height in meter square. BMI scores ≥30 kg/m^2^ were considered obese [17]. Central obesity was defined based on a waist circumference ≥102 or ≥88 cm for men and women, respectively [18].

#### Biochemical measurements

Twelve-hour fasting venous blood samples were collected for assessment of glycosylated hemoglobin (HbA1c), high density lipoprotein cholesterol (HDL-C), low density lipoprotein cholesterol (LDL-C), total cholesterol (TC), and triglycerides (TG) in accordance with the World Health Organization (WHO) guidelines.

#### Comorbidities

Major comorbid risk factors like hypertension, diabetes mellitus, and dyslipidemia were assessed. Criteria for diagnosis of hypertension were as per the Seventh report of the Joint National Committee on Prevention, Detection, Evaluation, and Treatment of High Blood Pressure (JNC7) [19]. In addition, the participants were considered to be hypertensive if they were previously diagnosed with hypertension or have been using any sort of anti-hypertensive medication regardless of their blood pressure readings. Diabetes mellitus was defined as per the WHO and American Diabetes Association (ADA) criteria as, HbA1c ≥6.5%, or by previous diagnosis of diabetes or were on anti-diabetes medication [20]. Dyslipidemia was diagnosed according to the WHO and the Third Adult Treatment Panel (ATP-III) of the National Cholesterol Education Program (NCEP) criteria. Dyslipidemia included, raised levels of TC, LDL-C, or TGs; and low levels of HDL-C, or if the subject reported using medications to lower blood lipid levels [21,22].

### Statistical analysis

Statistical analysis was performed by using SAS/STAT (SAS institute Inc. NC, USA). Continuous variables were presented as mean and standard deviation (SD), and categorical variables were summarized as number and percentage. Comparison between variables and significance testing was done using chi-square test or Fisher’s exact test or independent *t* test, as appropriate.

Multivariate logistic regression analysis was done to determine the factors influencing better awareness, with high awareness score as the dependent variable and other covariates including, age, presence of comorbid conditions and 5-year median length of stay as independent variables. A p value of <0.05 was considered statistically significant.

### Results

The study population consisted of non-Saudi employees working at a university and their family members (574 males and 306 females). The mean age (39.7 and 38.5 years) was similar for both male and female. The nationality of most of the study population was Middle East and North Africa (78%) while the rest were South Asians (22%). The study participants’ socio-demographic details, mean values of blood pressure, biochemical measurements, and prevalence of chronic diseases are summarized in **Table 1.** Most men attained higher levels of education and held academic positions in the university. Women had higher mean BMI than men (p<0.001). Gender differences were shown in men with higher mean systolic blood pressure (p=0.028), as well as TG and lower HDL-C values (p<0.001), posing risk of dyslipidemia; obesity was significantly higher among the females (p<0.001).

**Table 1.**
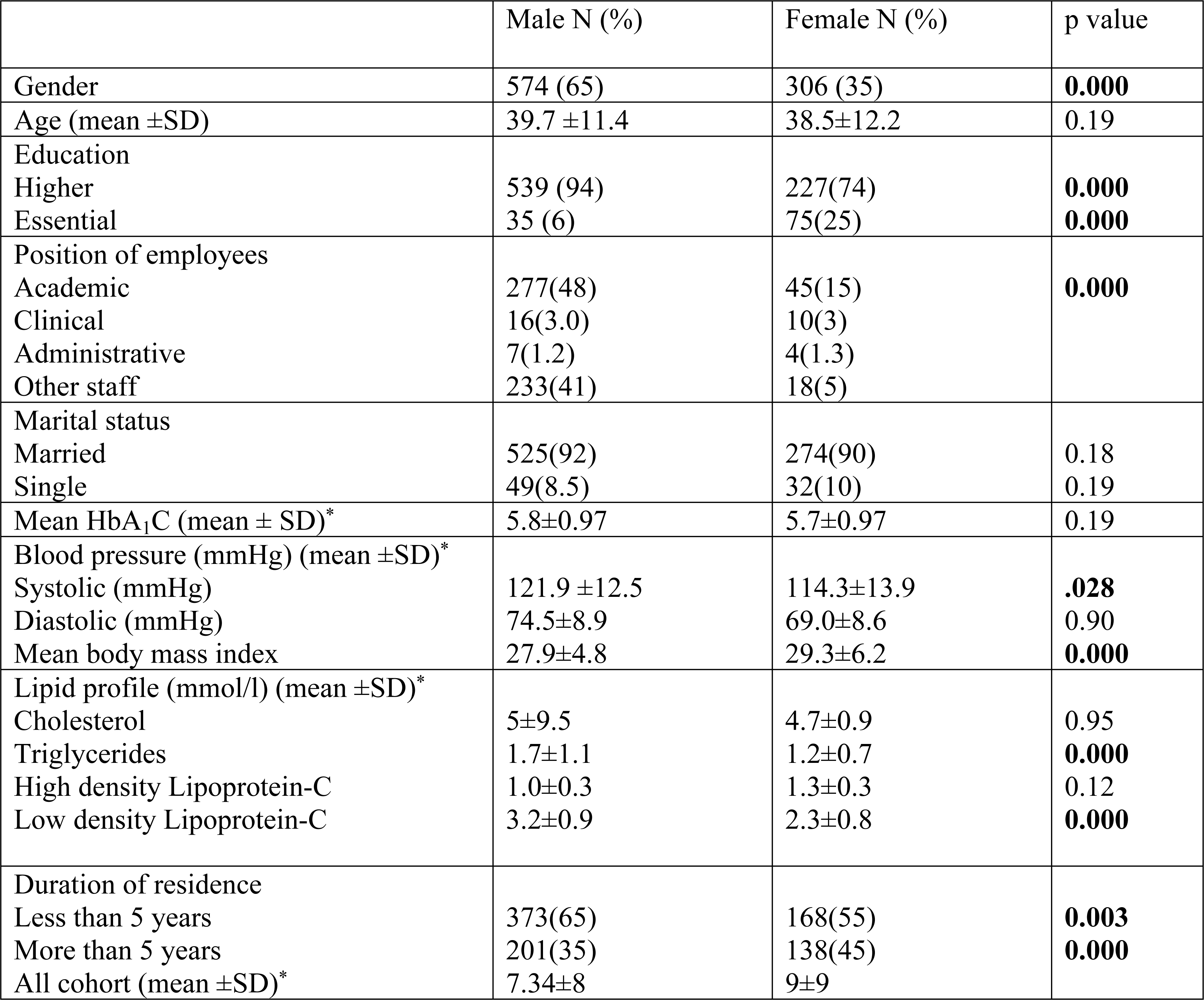
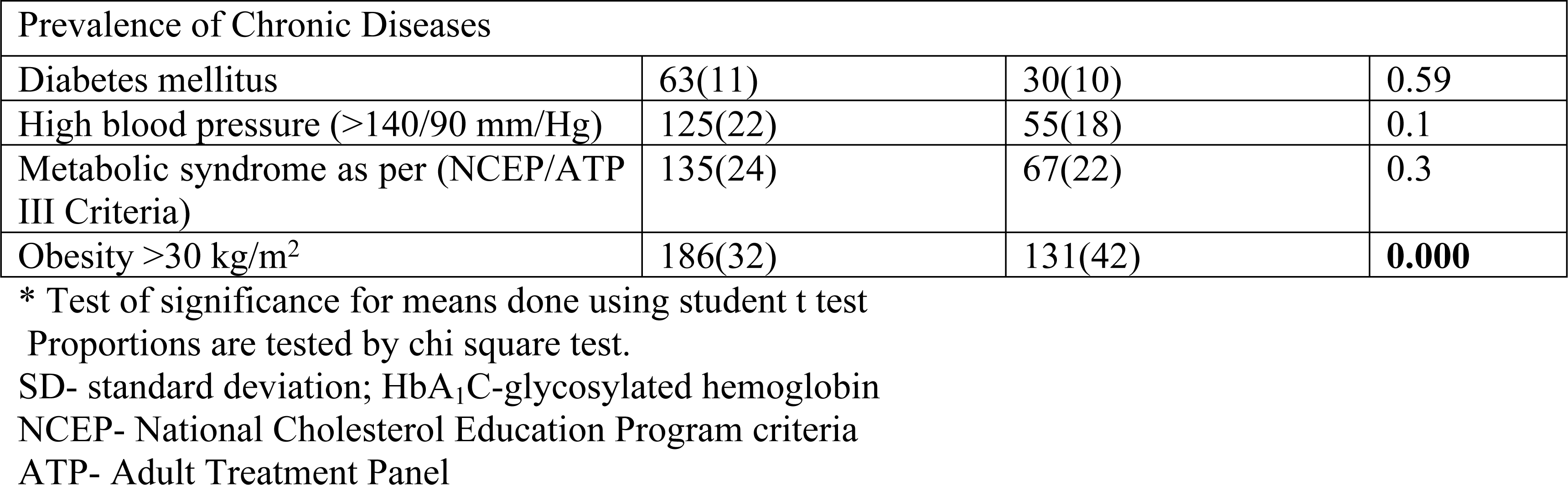
Details of demographic variables, biochemical measurements, and comorbidities of the study population:

The results of changes in the method of food preparation and dietary awareness in the overall population after migration are presented in f**igures 1 and 2** respectively. More than one third of the population adopted newer methods of food preparation. Barbeque was the most popular method, followed by the eating of raw vegetables (salads).Close to half of the study population showed substantial improvement in dietary awareness and exerted certain efforts to look for ingredients and find healthier options during food purchase.

**Figure 1:**
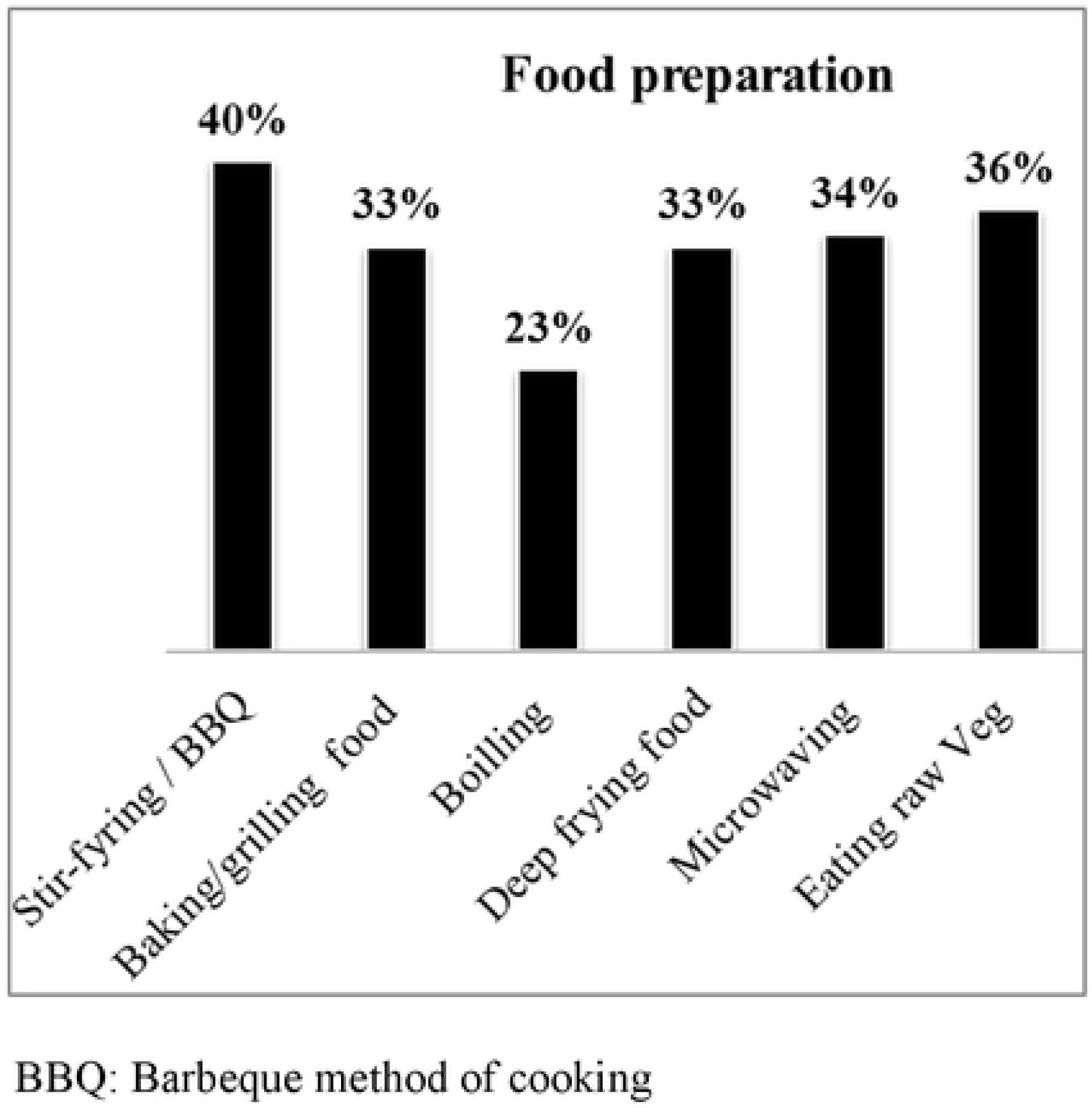
Chart depicting the reported changes in food preparation methods after residing in Saudi Arabia

**Figure 2:**
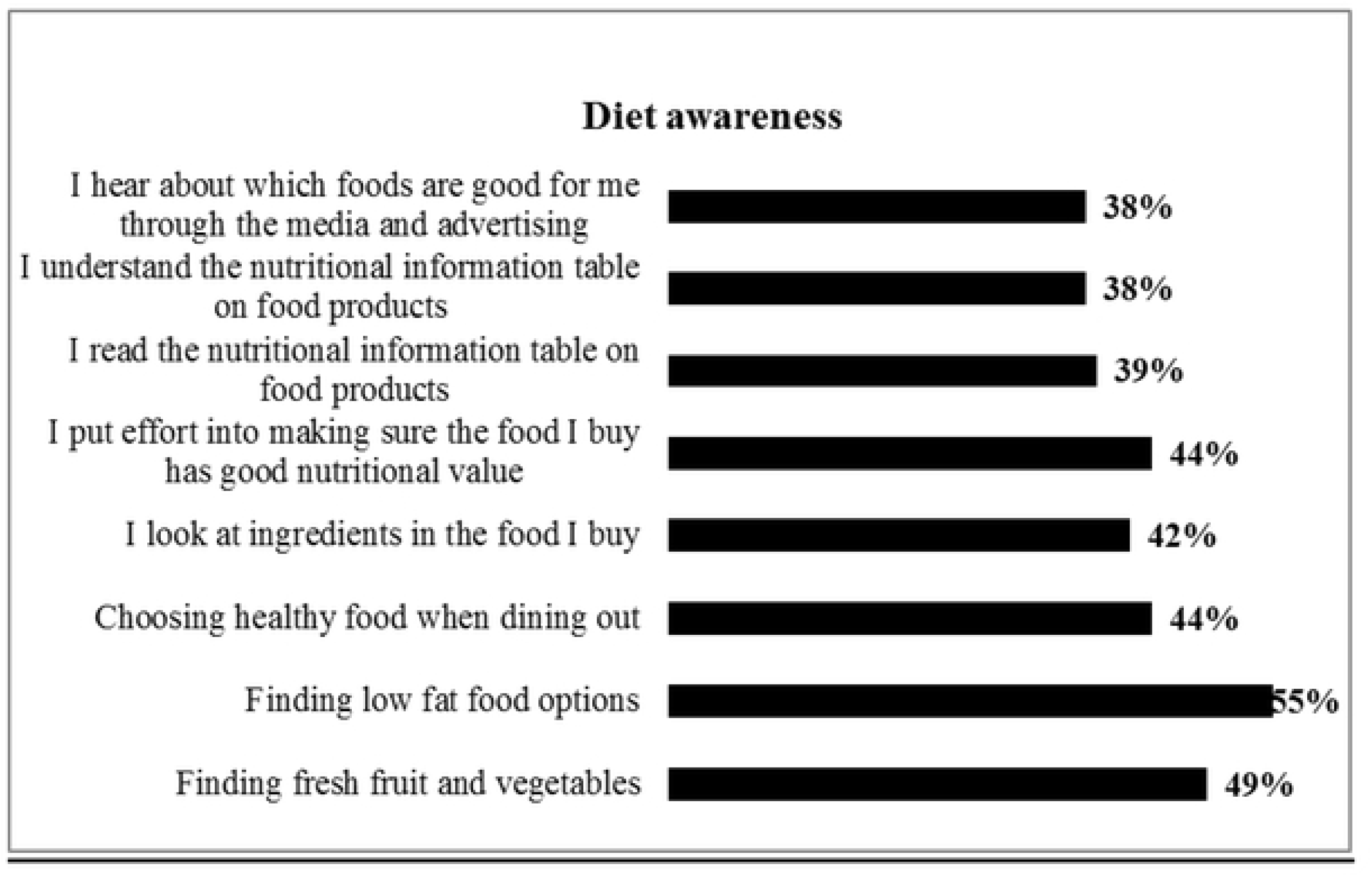
Percentages of the changes in diet awareness after moving to Saudi Arabia

**Table 2** summarizes the pattern of changes in food behavior due to dietary acculturation and chronic disease status among the participants, by duration of residency. Food behavior included portion size, method of food preparation, types of food consumed, and label reading behavior during food purchase. Size of food portion did not show any variation with migration. Those residing in Saudi Arabia for <5 years showed significant increase in the use of barbeque and stir-fry (p=0.018) as a method of food preparation while those with longer duration of residence status preferred cooking by microwave (p=0.002). There was an increase in salad consumption after migration (p<0.001). The other methods of food preparation like baking, boiling, and deep frying, did not show significant change after migration. Soft drinks and soda, high carbohydrate diet (potatoes and rice), high fat fried foods, and desserts and candies were consumed in significantly higher proportion among recent immigrants (<5 years residency). Label reading behavior and finding low fat options during purchase of food items was significantly higher among the ≥5 years residency duration group (p<0.001). The results showed higher prevalence of chronic diseases like diabetes (p<0.000), obesity (p<0.001), and blood pressure (p<0.012) in participants with longer residency status in Saudi Arabia.

**Table 2.**
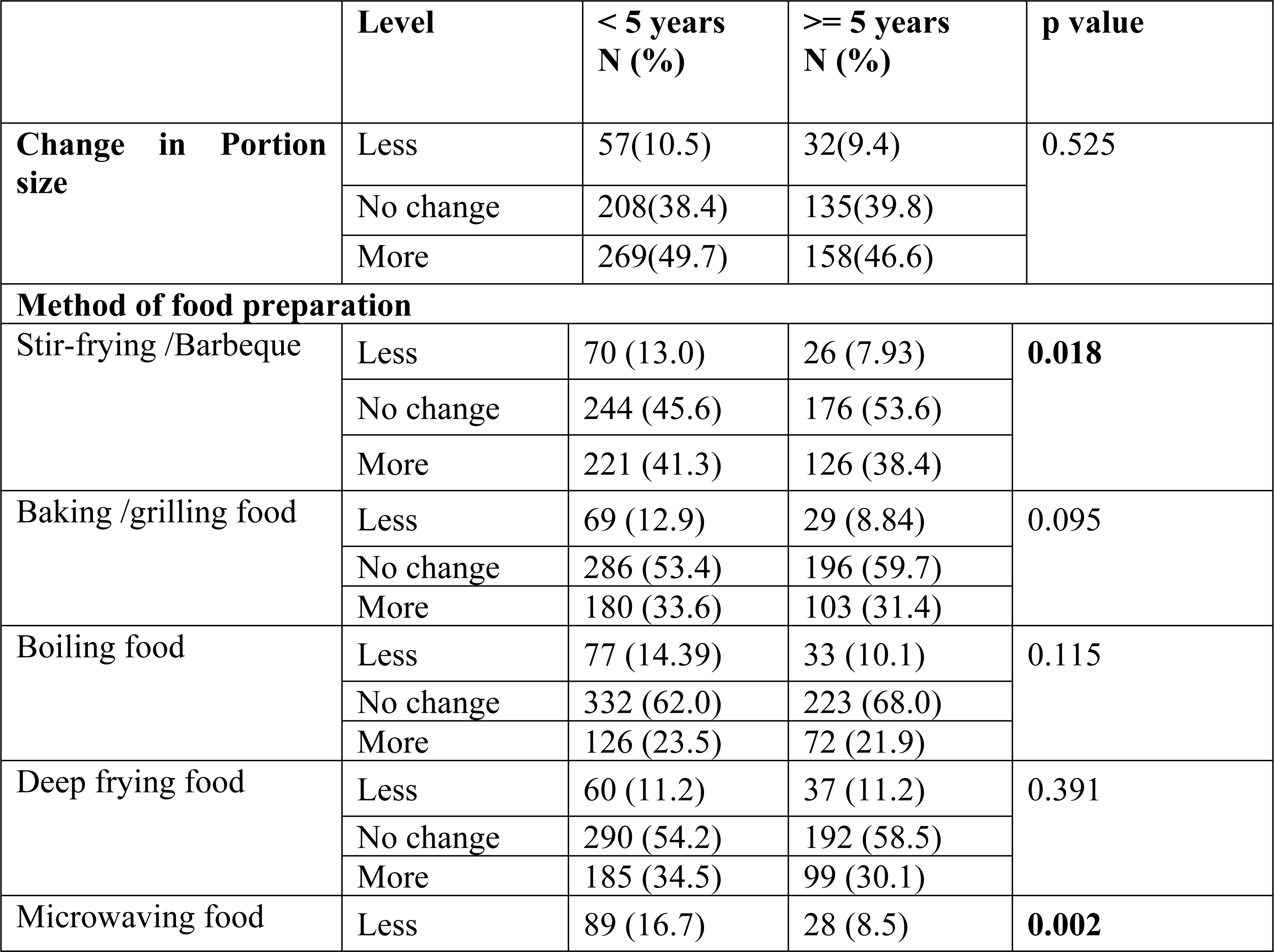
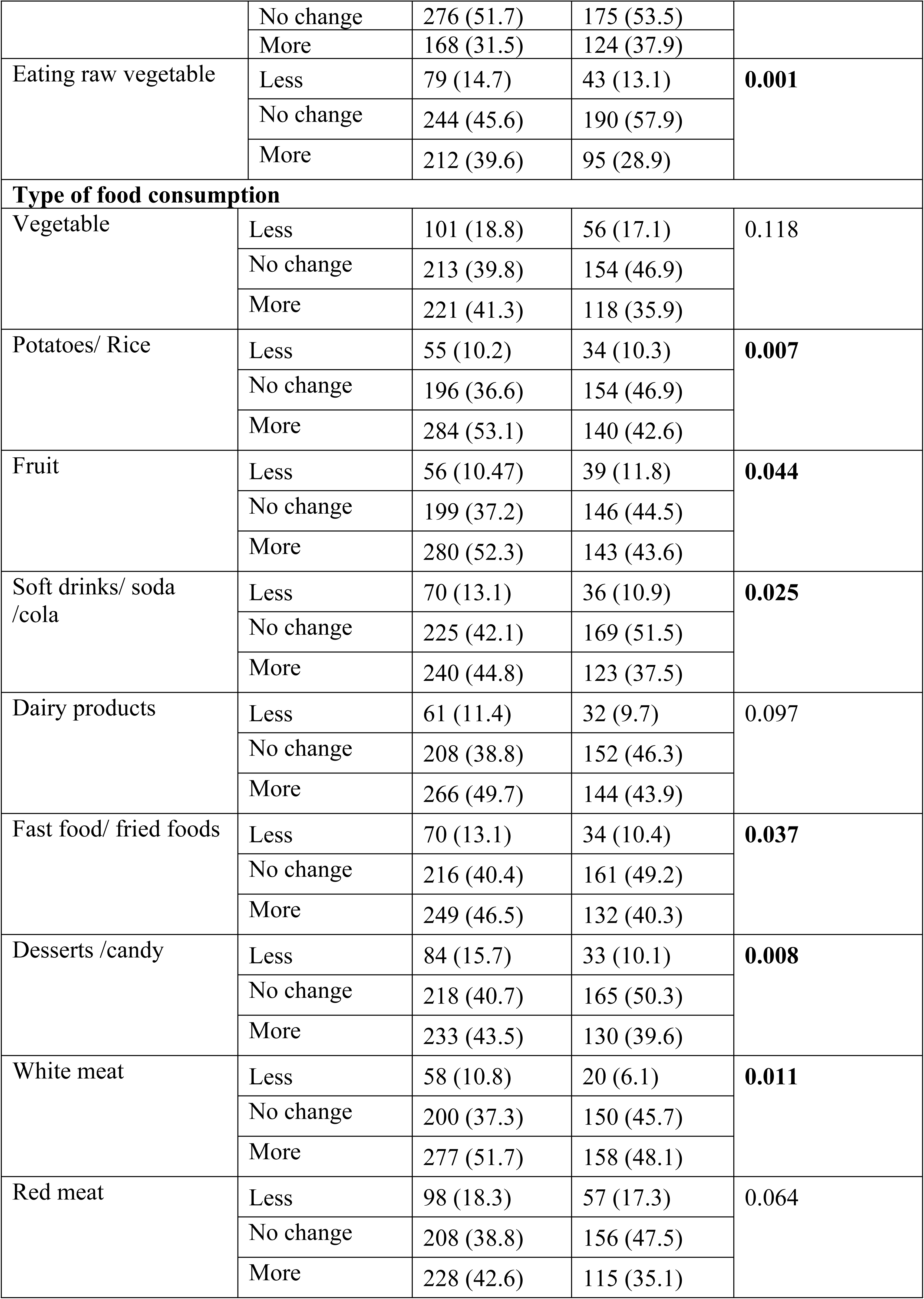
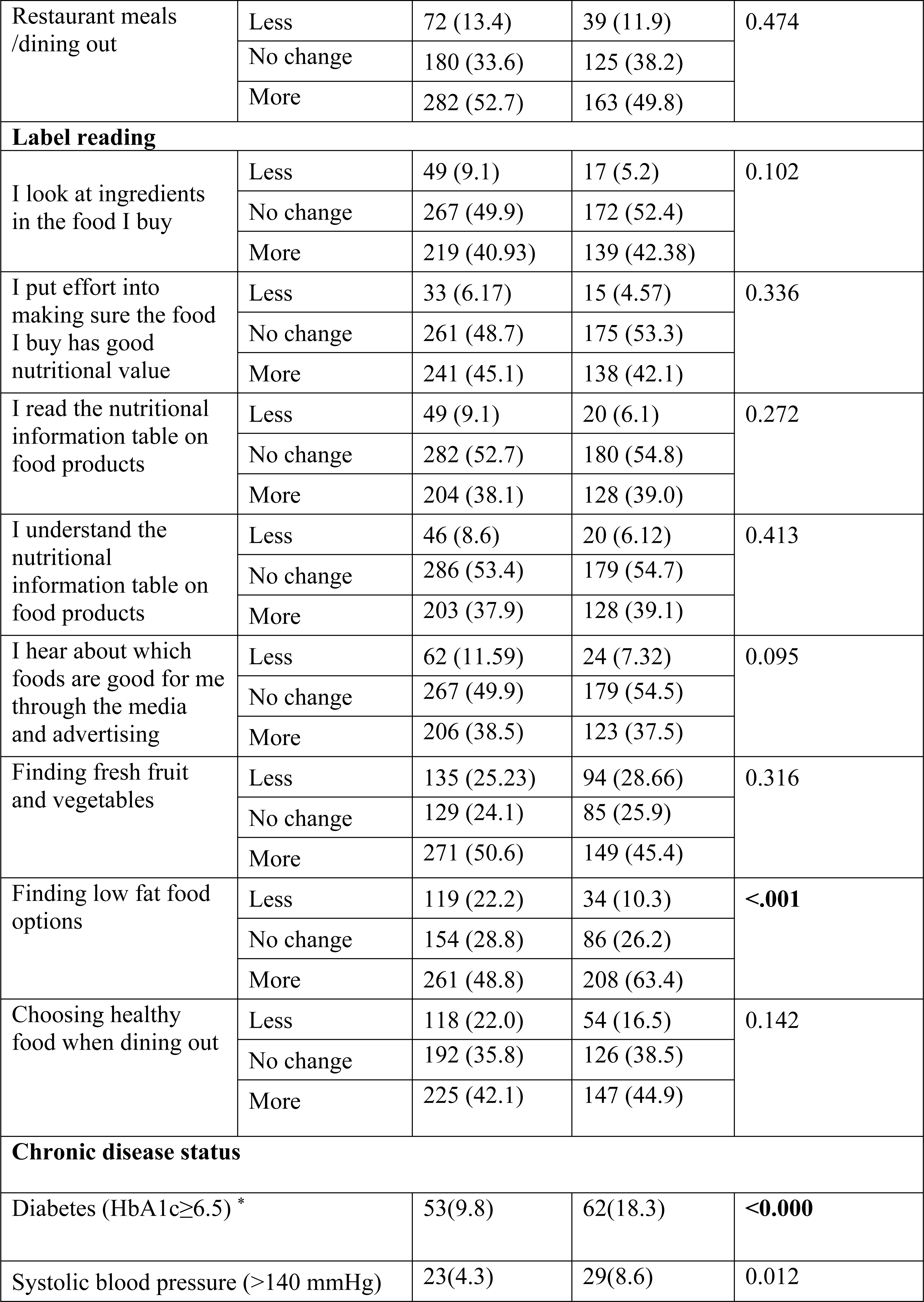
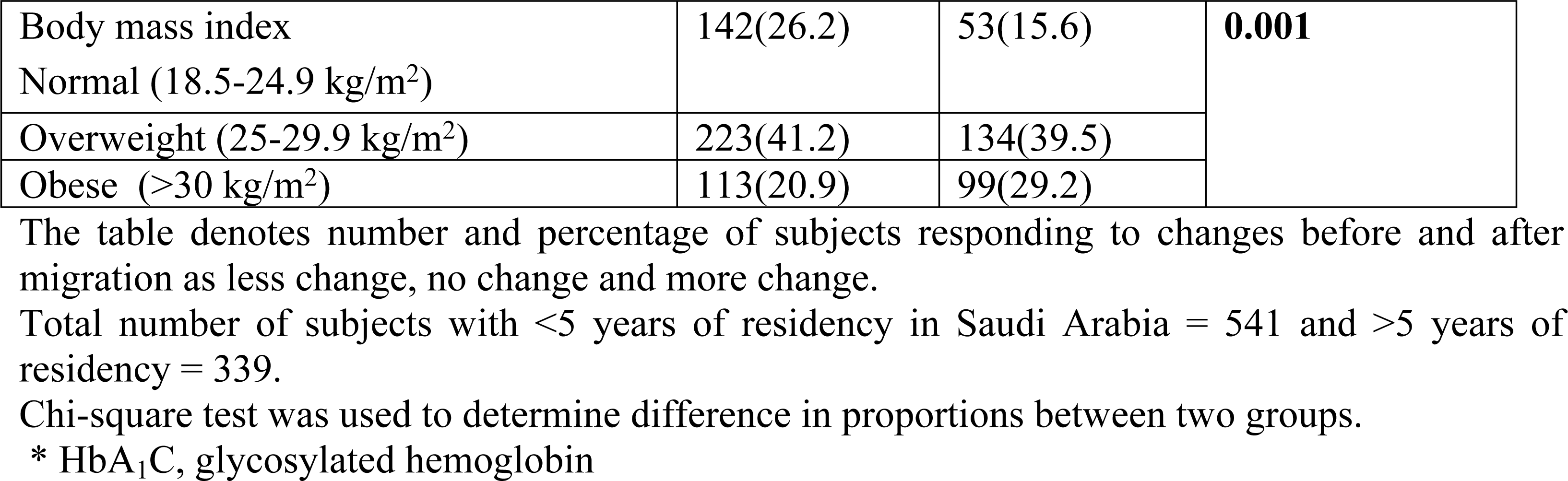
Changes in the method of food preparation, food consumption, food purchase behavior and change in chronic disease status by duration of residency in Saudi Arabia

**Table 3** describes the distribution of awareness and practice scores of healthy dietary habits. There were higher scores for both awareness (p=0.06) and practice (p<0.0001) of healthy food behavior in females than males. Longer duration of residency improved awareness (p=0.004) and presence of disease showed better practice score (p=0.003).

**Table 3.** Distribution of awareness and practice scores by gender, duration of residence status, and presence of comorbidities

| Variables | Awareness | Practice |
| --- | --- | --- |
| **Gender** |  |  |
| Male | 6.9±1.47 | 9.4±1.58 |
| Female | 7.1±1.33 | 9.8±1.39 |
| P value | **0.06** | **<0.001** |
| **Length of stay in Saudi Arabia** |  |  |
| Less than 5 years | 6.8±1.51 | 9.5±1.53 |
| Greater than or equal to 5 years | 7.14±1.27 | 9.6±1.52 |
| P value | **0.004** | 0.110 |
| **Comorbidity** |  |  |
| Overall | 6.9±1.43 | 9.5±1.53 |
| Normal, free of diseases | 6.9±1.44 | 9.4±1.60 |
| Presence of any disease | 7.1±1.39 | 9.7±1.35 |
| P value | 0.139 | **0.003** |

Maximum attainable awareness score = 8 and practice score = 14.
Awareness and practice scores reflect on the good and healthy practices related to dietary and food purchase behavior. No negative scoring was given for unhealthy dietary practices.
Comorbidity relates to the subject having one or more of the disorders such as diabetes, hypertension and/or dyslipidemia

**Table 4** shows the results of multivariate logistic regression analyses to determine the factors associated with higher awareness and practice scores. Presence of diabetes was the only independent variable found to be significantly associated with higher awareness. Those with diabetes were two times more aware of healthy food than those without diabetes (odds ratio [OR] = 2; confidence interval [CI]= 1.2-3.2; p=0.005). And it was interesting to note that practice was not influenced by other covariates like age, gender, length of stay and presence of comorbidities.

**Table 4.**
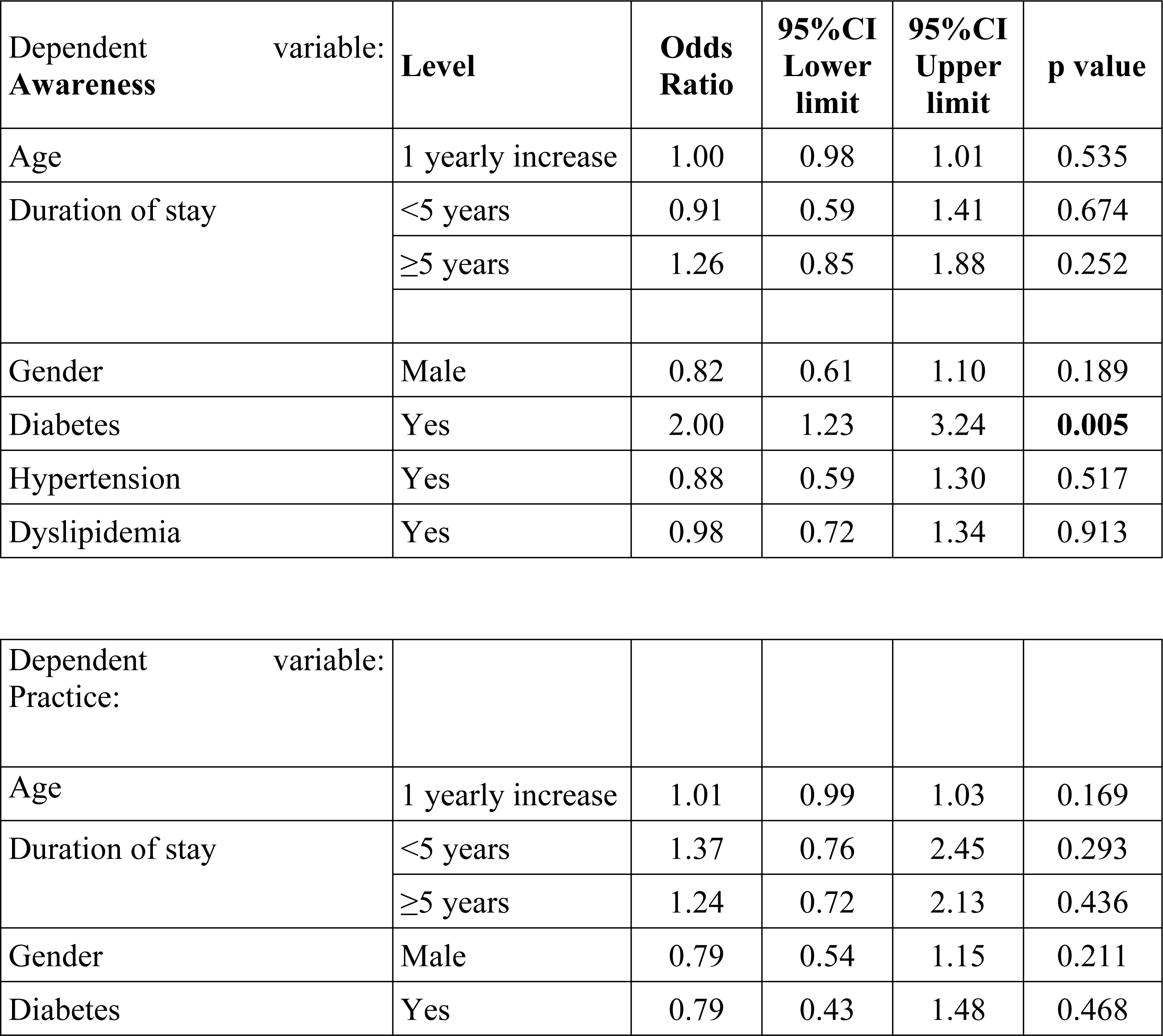
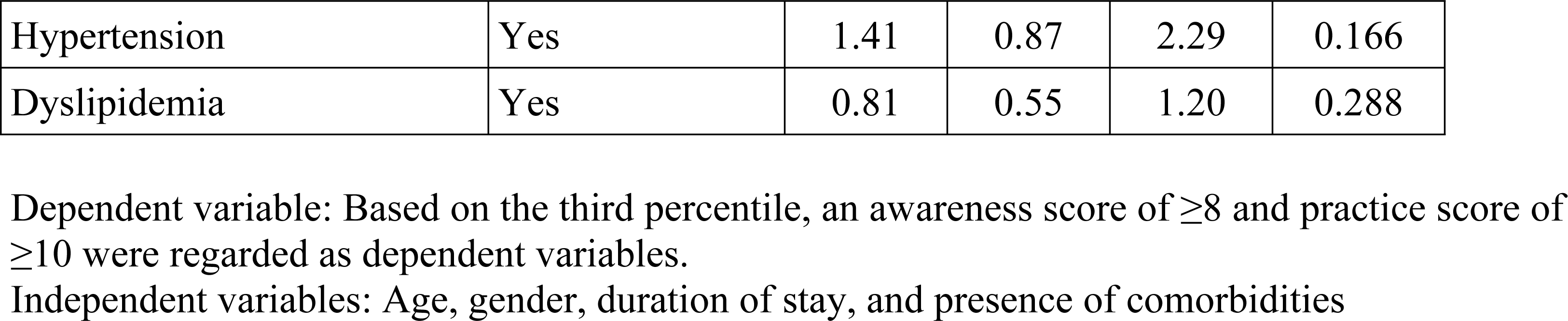
Multivariate logistic regression analysis of factors associated with higher awareness and practice scores

Changes in method of food preparation due to migration in the presence and absence of comorbidities were also assessed and presented in **Table 5**. The study participants with comorbidities mostly preferred not to change their previous method of food preparation.

**Table 5.** Level of changes in the method of food preparation before and after migration by the presence or absence of comorbid conditions

| Covariate | Level | Comorbidities<br>Present<br>N=293 | Absent<br>N=587 | p value |
| --- | --- | --- | --- | --- |
| Stir-frying /Barbeque | Less often | 24 (8.54) | 72 (12.37) | **0.05** |
|  | No change | 152 (54.09) | 268 (46.05) |  |
|  | More often | 105 (37.37) | 242 (41.58) |  |
| Baking /grilling food | Less often | 22 (7.83) | 76 (13.06) | **0.04** |
|  | No change | 170 (60.5) | 312 (53.61) |  |
|  | More often | 89 (31.67) | 194 (33.33) |  |
| Boiling food | Less often | 29 (10.32) | 81 (13.92) | 0.30 |
|  | No change | 188 (66.9) | 367 (63.06) |  |
|  | More often | 64 (22.78) | 134 (23.02) |  |
| Deep frying food | Less often | 31 (11.03) | 66 (11.34) | 0.65 |
|  | No change | 163 (58.01) | 319 (54.81) |  |
|  | More often | 87 (30.96) | 197 (33.85) |  |
| Microwaving food | Less often | 38 (13.57) | 79 (13.62) | 0.95 |
|  | No change | 145 (51.79) | 306 (52.76) |  |
|  | More often | 97 (34.64) | 195 (33.62) |  |
| Eating raw vegetable | Less often | 25 (8.9) | 96 (16.49) | **0.007** |

| Covariate | Level | Comorbidities Present<br>N=293 | Absent<br>N=587 | p value |
| --- | --- | --- | --- | --- |
|  | No change | 160 (56.94) | 274 (47.08) |  |
|  | More often | 96 (34.16) | 211 (36.25) |  |

## Discussion

### Dietary acculturation by duration of residency

The results indicate bidirectional findings. The effect of dietary acculturation had certain favorable influence with improvements in healthy food behavior in addition to acquisition of negative dietary habits as well. Our participants showed marked improvements in the practice of oil-free methods of cooking like barbeque, baking and microwaving, consumption of raw food and improved label reading behavior, and search for low fat options during food purchase, which can be regarded as the manifestation of positive dietary acculturation after migration into Saudi Arabia. However, at the same time, high consumption of fast food, carbonated drinks, and sugared food demonstrated a significant negative consequence. The present study also reported an increase in the consumption of fats in the form of meat, fried food, and increased intake of potatoes and rice. Although awareness of healthy food improved substantially with migration, it did not necessarily translate into practice. However, the dietary habits gradually improved the longer the duration of stay.

These findings are consistent with a study similar to ours, conducted by Tiedje et al. that included a diverse group of immigrant population, such as Sudanese, Somali, Mexican, and Cambodian communities living in the United States (US), reported improved awareness in healthy food after migrating to the US but failed to correlate with healthy practices. With Americanization, they gradually adapted to the improved dietary practices [23]. Given the low practice scores compared to the awareness scores, our study also demonstrated a weak correlation between awareness and practice, indicating that improved knowledge does not necessarily reflect on healthy practices among recent immigrants. This is an issue of concern since the findings have implications on the health status of immigrants over time.

Many studies have documented mixed findings, due to the diverse nature of human adaptability, from traditional to modern food, depending on age and duration of residency [24,25]. Migration to affluent countries enable the immigrants to encounter gradual transition influenced by cultural and social acceptability of the host’s environment. Qualitative analysis of Arab immigrants to western nations has reported greater nutritional awareness with simultaneous inclination towards soft drinks and fast food [26]. Wander et al. demonstrated stage-wise pattern of changes in relation to duration of stay of Asian community in Oslo, reporting increased fat intake in the form of oil and meat initially, followed by reduced consumption over time [27]. Lesser et al. studied the dietary behavior of South Asians (India, Pakistan, Bangladesh, and Sri Lanka) after migration to western nations and found favorable changes similar to that in our study, in healthier methods of food preparation with a simultaneous increase in carbonated drinks, fried food, and convenience food [6]. These findings are suggestive of a similar pattern of adaptation following migration to Western nations. Saudi Arabia, in addition to the availability of traditional food, can also be considered to be highly westernized in terms of food pattern and easy availability of ready-to-eat, processed, and fast food as in any other western nation. The newer immigrants in our study were at a greater risk of unhealthy dietary acculturation, while the older immigrants with longer duration of stay had better nutritional awareness in determining food preferences and healthy dietary choices.

Increased portion of the meal size, in about half of the surveyed population, was reported after migration. Likewise, increased frequency of dining outside, although insignificant in the present study, still has important implication, since more than half of the participants frequently dined out. The need for standardized portion sizes in restaurants has always been emphasized by researchers and by the medical community since people tend to consume excess calories while dining out. Cohen and Mary published a report on commercial fast food giants serving calories ranging from 785 to 1860 per ordered meal per person; specifying an unambiguous surplus of calorie intake, which poses risk for chronic diseases [28]. Of utmost concern, these findings signal the hidden risk, underlying increased consumption of food. At the outset, it is worthwhile to mention an interesting policy proposal by the Government of Canada’s anti-obesity plan where proposals are being developed to consider shrinking the size of a standard pizza serving to not more than 928 calories in a desperate attempt to tackle the rising obesity rates among the Canadian native and immigrant population [29]. Such initiatives may prove beneficial in cutting down calories by the method of forced implementation rather than by mere health warnings in the form of awareness campaigns.

### Chronic disease status and dietary acculturation

It is well documented and well-reviewed that new immigrants enjoy better health status than the natives, which has been termed as ‘healthy immigration effect,[30] but the decline in health status with the passage of time has been strongly linked to challenges in dietary acculturation and stress [31,32]. In addition, unhealthy dietary lifestyle has strongly been associated with chronic diseases as a major risk factor [33]. Our finding of high prevalence of all the three major chronic diseases; diabetes, hypertension, and obesity among immigrants with longer residency, is a matter of concern. This finding must however reflect on better awareness and practices, but the multivariate analysis showed that only diabetes, among all the comorbidities, had an influence on awareness of healthy diet. Furthermore, healthy practices did not appear to be associated with the presence of any of the diseases. The possible explanation for this finding could be the efforts of the national diabetes control program, which prioritizes mass education on diabetes prevention. The practices remain an individual’s choice and depend on the individual’s personal perception and attitude towards health. The present study showed an inclination towards fast food after migration. These findings certainly have substantial implications on health, prompting necessary action by policy makers. By increased affordability and vast availability of almost every kind of commercial giants selling fast foods in Saudi Arabia, not only the immigrants but the population *en masse* is at risk of obesity and other cardiovascular disorders.

### Limitations of the study

Generalizability of the results of this study is one of the major limitations. Since the study participants were enrolled from a single center, the study sample may not be representative of the population. Moreover, the study population was well-qualified in terms of education; as a result, the role of literacy in dietary acculturation could not be assessed due to the absence of less educated groups in the sample. Additionally, the high level of education would have also contributed to the increased levels of awareness. We therefore recommend future research involving a large representative sample of the population. In the present study, the cross sectional study design used could not establish causation and could only report the frequency at one point in time. Cohort study is highly recommended to determine the process of changes in dietary pattern and health status of the immigrants. The present study was not a Knowledge-Attitude-Practice study; hence, it was not intended to measure the attitude of the immigrants towards dietary behavior. However, to our knowledge, this is the first study in the region of Saudi Arabia to have examined the effects of migration on diet behavior during health and disease. The results could still be considered to have certain significant implications on diet-related risk behavior and to serve as an important source of data for future research.

## Conclusion

Migration into Saudi Arabia showed marked changes in methods of food preparation and food choices, but acquisition of unhealthy dietary practices also co-existed despite improved awareness and despite the presence of comorbidities. These findings suggest the need to conduct population-based studies, involving multi-ethnic communities to provide evidence that could be used by policy makers to ensure standardization of dietary regulations of certain foods, to limit immigrants’ fat intake and thereby reduce the risk factors of non-communicable diseases.

## Authors contribution

RA Conceptualized, conducted the study, ST performed part of analysis and wrote the manuscript, AH supervised the study, AU performed the analysis

## Acknowledgement

We are thankful to all the participants of the study. The authors are grateful to the Deanship of Scientific Research, King Saud University for funding through the Vice Deanship of Scientific Research Chairs. Furthermore, authors thank the Deanship of Scientific Research and RSSU at King Saud University for their technical support.

